# Selective enhancement of low-gamma activity by tACS improves phonemic processing and reading accuracy in dyslexia

**DOI:** 10.1101/2020.04.15.042770

**Authors:** Silvia Marchesotti, Johanna Nicolle, Isabelle Merlet, Luc H. Arnal, John P. Donoghue, Anne-Lise Giraud

**Affiliations:** Department of Neuroscience, University of Geneva, Geneva, Switzerland; INSERM U1099, LTSI, Campus de Beaulieu, Rennes, France; Université de Rennes 1, LTSI, Campus de Beaulieu, Rennes, France; Brown University, Providence, United States

## Abstract

The phonological deficit in dyslexia is associated with altered low-gamma oscillatory function in left auditory cortex, but a causal relationship between oscillatory function and phonemic processing has never been established. After confirming a deficit at 30 Hz with electroencephalography (EEG), we applied 20 minutes of transcranial alternating current stimulation (tACS) to transiently restore this activity in adults with dyslexia. The intervention significantly improved phonological processing and reading accuracy as measured immediately after tACS. The effect was selective to 30 Hz stimulation, and proportional to dyslexia severity. Importantly, we observed that the focal intervention on the left auditory cortex also decreased 30 Hz activity in the right superior temporal cortex, resulting in reinstating a left dominance for the oscillatory response, as present in controls. These findings formally establish a causal role of neural oscillations in phonological processing, and offer solid neurophysiological grounds for a potential correction of low-gamma anomalies, and for alleviating of the phonological deficit in dyslexia.

## Main Text

Dyslexia is a frequent disorder of reading acquisition affecting up to 7% of schoolchildren, and characterized by persisting difficulties with written material throughout adulthood. Identifying the neural bases of dyslexia to devise efficient treatments has motivated intense research in the last decades [1–3]. These treatments include behavioral auditory and reading training [4,5] and more recently non-invasive electrical brain stimulation [6–8], even though the exact underlying action mechanisms remain uncertain. From a neuroscience viewpoint, dyslexia poses an interesting challenge because it selectively affects one aspect of language, the mapping of phonemes onto graphemes, while leaving other cognitive domains intact, such as speech perception and production, or visual and auditory processing [9]. Although several possible causes have been proposed for dyslexia [10], the predominant one is a phonological deficit, i.e. a difficulty to process the sounds of language. The deficit primarily affects phonological awareness, conscious access, representation and internal monitoring of speech sounds [11–13], the capacity to form rich categorical phonemic representations [13,14], as well as naming and verbal memory [15].

Unlike oral language that arises through mere exposure, reading requires explicit learning, through which children become aware that the syllables they are used to parse (e.g. via nursery rhymes) are made of smaller units, the phonemes. Learning to read consists in mapping these new basic phonological building blocks to specific visual symbols. This so called phoneme/grapheme mapping is only possible if the child is able to match the sound associated with the visual symbol with his/her own phonemic inventory, made of infra-syllabic elements that can be taken out and replaced by another articulable sound [16]. At the acoustic level, critical phonemic contrasts (e.g. /t/ versus /d/ or /da versus /b/, etc.) are underpinned by rapid events (noise bursts, formant transitions, voicing etc.), and grasping them requires auditory sampling at a rate allowing for their neural encoding as individual patterns. The optimal sampling rate should hence enable the integration of basic 30-40 ms acoustic segments, comprising roughly vocal tract occlusion and peri-occlusion cues such as voicing [17].

Theoretical models propose that neural oscillations in the 25-35 Hz range could be the basic speech sampling rate, from which would derive (in an evolutionary sense) the inferior bound of the phonemic temporal format, i.e. the shortest possible linguistic unit (∼25-30 ms) that can be individually represented [18]. Basic neurophysiological studies confirm that two sounds must be separated by at least 25 ms to be perceptually and neurally individualized [19]. In line with these observations, neuroimaging studies have repeatedly linked dyslexia with a deficit in oscillatory activity in the low-gamma band [20–24]. This deficit could be related to difficulties in the processing of rise-time in amplitude modulated sounds, typically associated with anomalies in slower neural oscillation ranges [24–26]. Young adults with dyslexia show disrupted low-gamma 30-Hz response in left auditory cortex, and abnormally strong responses at higher frequencies (around 40Hz), suggesting that auditory sampling could be faster than in typical readers [22,27]. This double anomaly has been associated with a deficit in respectively phonological processing (due to atypical phonemic format) and working memory (due to more basic phonemic elements per language unit to hold in memory, [22]). Furthermore, consistent with atypical morphological asymmetry of temporal cortex in dyslexia [28], left-dominance of the low-gamma auditory response is also disturbed. In both cases (anatomical and functional) the asymmetry anomaly is statistically related with the phonological deficit [22].

These findings converge toward a possible oscillation theory of phonemic construction and of the phonological deficit in dyslexia, originating primarily in atypical functioning of the left auditory cortex. This theory is worthy of investigation as it could offer an easy entry point for therapeutic interventions through non-invasive neuromodulation aiming at normalizing oscillatory function in auditory cortex [29]. Building on the hypothesis that disrupted low-gamma activity in left auditory cortex could be causally related to the phonological deficit in dyslexia, we tested whether restoring low-gamma oscillatory function in individuals with dyslexia could improve phonemic perception and indirectly reading performance.

To address this question, we used high definition transcranial alternating current stimulation (tACS) with a 4×1 ring electrode configuration, which was focal, painless, frequency-specific, under the assumption that stimulating at 30 Hz should boost neural activity at the same frequency [30,31]. We conducted this neuromodulation study in a single-blind way in thirty adults (15 with dyslexia, 15 fluent readers, mean age 26.4 years, SD ± 8.1, range 18-47) involving for each subject 22h of experimental procedures spread over 4 experimental days (Fig **S1**). After a thorough assessment of dyslexia in all participants on Day 1, they performed on Days 2-4 a custom-designed battery of linguistic tests (Table **S1**) probing reading efficiency (speed and accuracy) and phonological processing (via pseudo-word test i.e. non-lexical word repetition, and spoonerism i.e. inverting the first phoneme of a word pair). Prior to the linguistic-tests battery, we recorded auditory-steady state response (ASSR) to pure tones modulated in amplitude with a fixed frequency (range: 28-62 Hz) through a 64 channels electroencephalography (EEG) system. EEG data were also acquired prior and after tACS intervention in order to confirm reduced 30 Hz responses in individuals with dyslexia relative to normo-readers, and monitor the neurophysiological changes subsequently induced by tACS.

ASSRs and behavioral tests were performed *before, immediately after* and *one hour after* 20 minutes of focal tACS (<= 2mA stimulation) over left auditory cortex, a stimulation duration that is argued to be effective while minimizing potential side effects [32]. The specificity of the 30 Hz intervention was controlled using two within-subject conditions: an active control stimulation at 60 Hz probing whether the expected effects could be merely due to delivering alternating current, and a placebo stimulation (sham condition) where no current was delivered. Given the selective deficit in left auditory cortex in subjects with dyslexia, we expect focal tACS to partly restore the 30 Hz neural response in that specific region in the dyslexia group inducing a benefit on phonemic processing, whereas no such effect is expected after 60 Hz stimulation in either group.

## Results

Before examining the effect of tACS on phonemic processing and reading, we first tested whether we could reproduce the 30 Hz deficit in the left auditory cortex that was previously shown in subjects with dyslexia [20,22,27]. This finding was a requirement to address whether tACS could effectively normalize neural activity. We analyzed the EEG auditory responses to amplitude-modulated sounds at frequencies between 28 and 62 Hz as measured *before* tACS. Auditory stimuli evoked typical waveforms with N100 and P200 auditory components [33] and topographies, with the strongest peak-to-peak response amplitude at FCz (Fig **1A**). In a first approach, we restricted the EEG analysis to this electrode. In line with a large body of literature [34,35], the time-frequency transformation of the EEG signal during the steady-state interval (auditory steady-state response, ASSR) peaked for auditory stimuli delivered with a 40-Hz amplitude modulation Fig **3A**). As the 30 Hz response deficit in dyslexia was expected to be lateralized and to dominate in *left* auditory cortex [22], we also reconstructed the activity of neural generators responsible for the signal recorded on the scalp. Based on previous results [28,36], we performed a source space analysis on two regions of interest (ROIs) in each cerebral hemisphere, corresponding to auditory cortex and superior temporal gyrus. Source reconstruction obtained from scalp recording also identified the strongest responses at these locations (Fig **S4**). We confirmed a statistically significant deficit in 30 Hz activity in the left auditory cortex in subjects with dyslexia relative to controls (T_27_ = 2.1, p < 0.05, d = 0.77, Fig **1C****-left**). We also confirmed that the effect was not present in the right auditory cortex (T_27_ = 0.068, p > 0.05, d = 0.02, Fig **1C****-right**), indicating that the deficit was lateralized. Further, there was no 30 Hz deficit in the superior temporal region surrounding auditory cortex in either hemisphere, confirming previous observations [22].

**Fig. 1.**
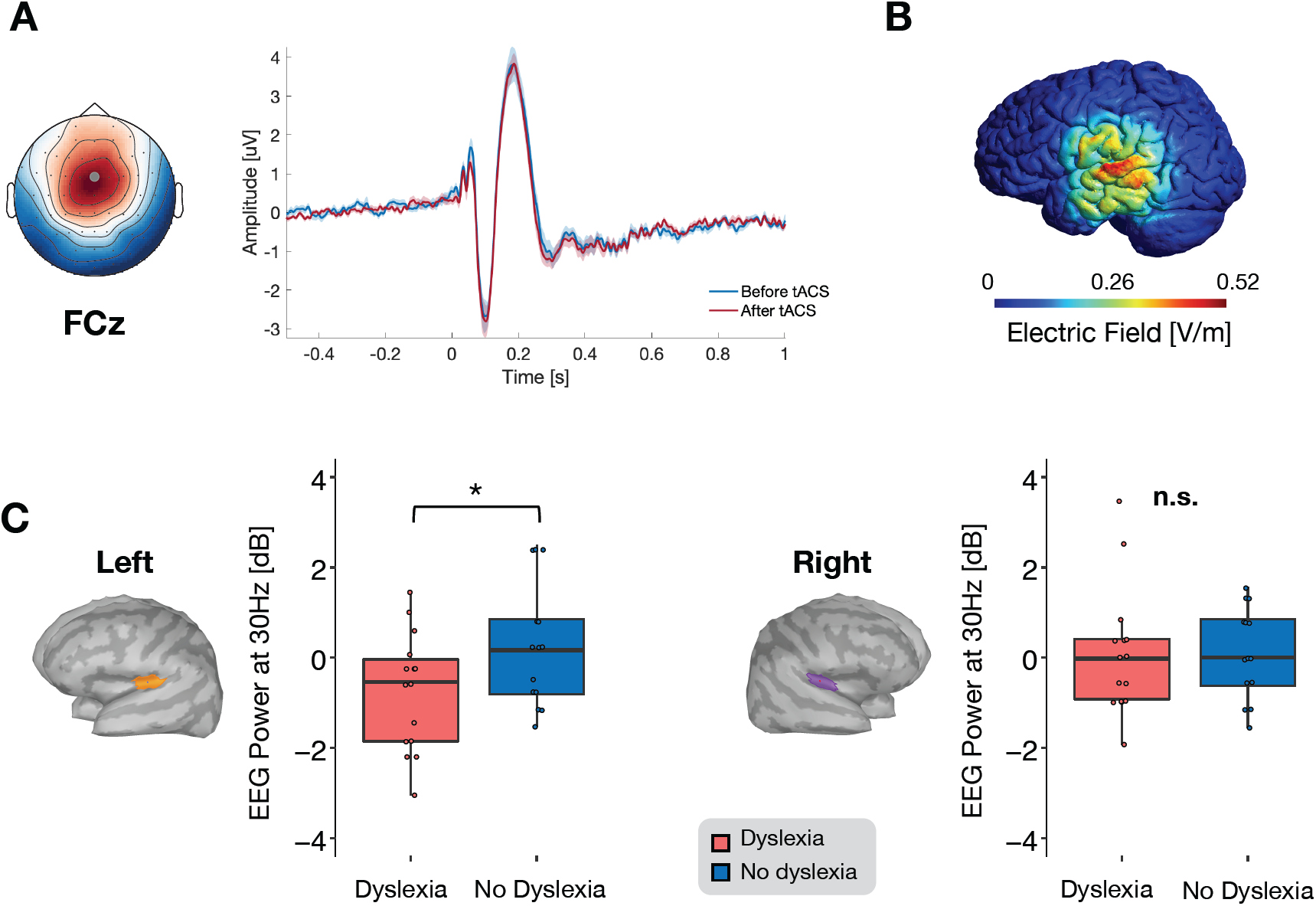
Auditory evoked response at the level of scalp electrodes, tACS induced electric field reconstruction and group difference at baseline at 30 Hz. (**A**) The average response over time (dyslexia group, all AM frequency considered) showed the stereotypical auditory N100 and P200 components and related topographies. The strongest peak-to-peak response amplitude was recorded at electrode FCz for both the N100 and P200 components (**B**). Simulation of the electric field induced by the tACS stimulation using a high-definition 4×1 electrodes configuration (model obtain with the freeware softwares SIMNIBs, www.simnibs.de and *gmsh*, www.gmsh.info. (**C**) Source reconstruction of the response to 30 Hz AM at 30 Hz recorded in the session *before* the 30 Hz tACS. In the left auditory cortex, the dyslexia group showed a reduced response as compared to the no dyslexia group (left); no difference between the groups was found in the right hemisphere (right).

**Fig. 2.**
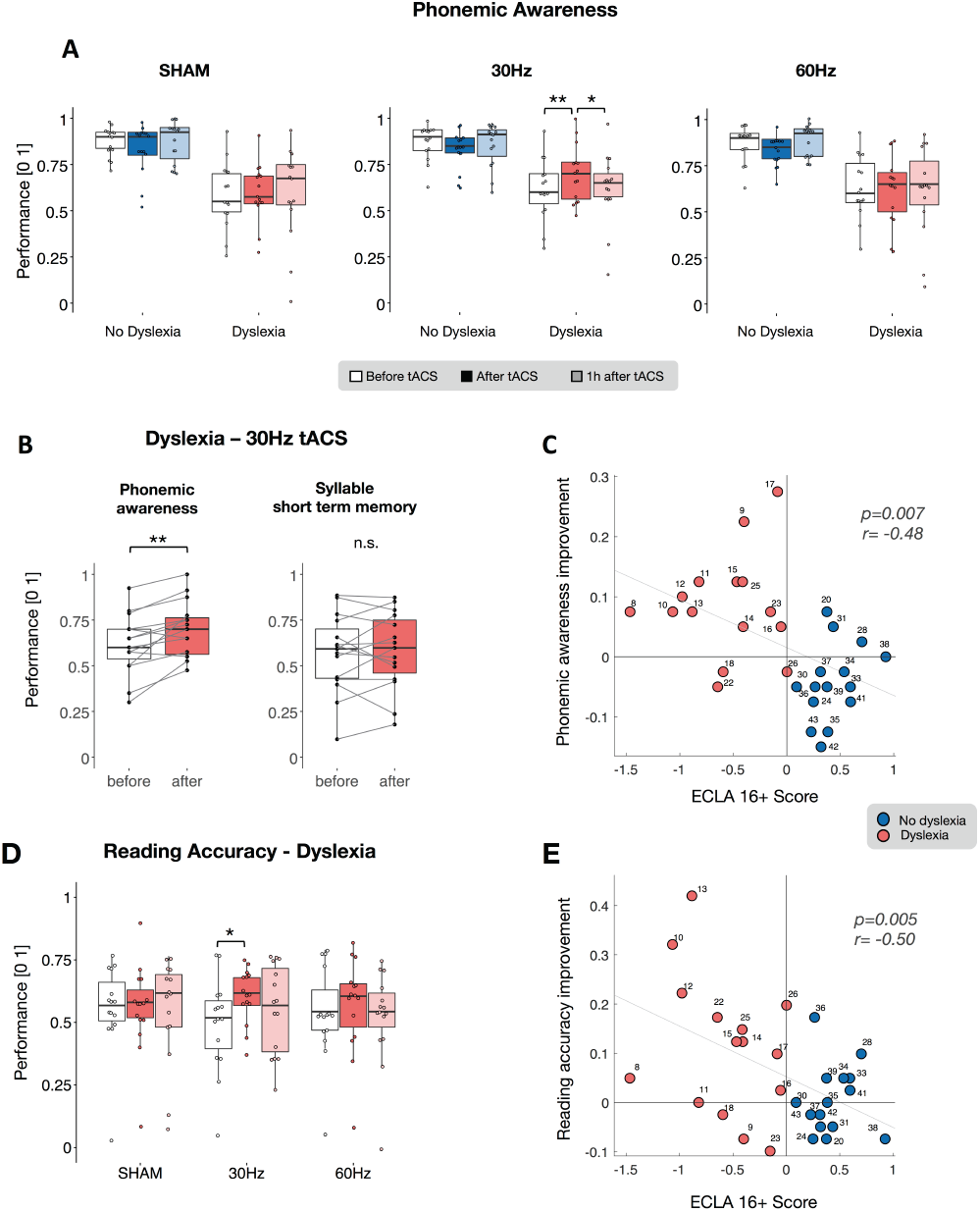
tACS stimulation effectiveness: behavioural results. Phonemic awareness for each tACS condition (sham, 30 Hz and 60 Hz) and for the sessions *before* (white bar plot), *after (*dark color), and *one hour after tACS* (light color) in the dyslexia (salmon) and no dyslexia (blue) group (**A**). This index was obtained by considering the number of errors in the repeated phonemes at the pseudo-word test (z-score values). Statistical analysis showed an interaction between group and session only for the 30 Hz stimulation: the dyslexia group showed performance improvement *after* the 30 Hz tACS as compared to the session *before* the stimulation, and decreased back to baseline values *one hour after* the tACS (* for p < 0.05, ** for p < 0.01). For the same tACS condition, performance of participants without dyslexia did not statistically change across sessions. The improvement in phonemic awareness *immediately after* the 30 Hz tACS in the dyslexic group (**B**, left) was not accompanied by changes in syllable short term memory (**B**, right). The degree of improvement in phonemic awareness following the 30 Hz tACS expressed a strong negative relationship with linguistic skills measured with the ECLA16+ test (**C**, negative values correspond to poorer linguistic skills). Reading accuracy improved *after* the stimulation at 30 Hz as compared to the session *before* **(D)**. This improvement correlated negatively with the ECLA16+ **(E)**.

**Fig. 3.**
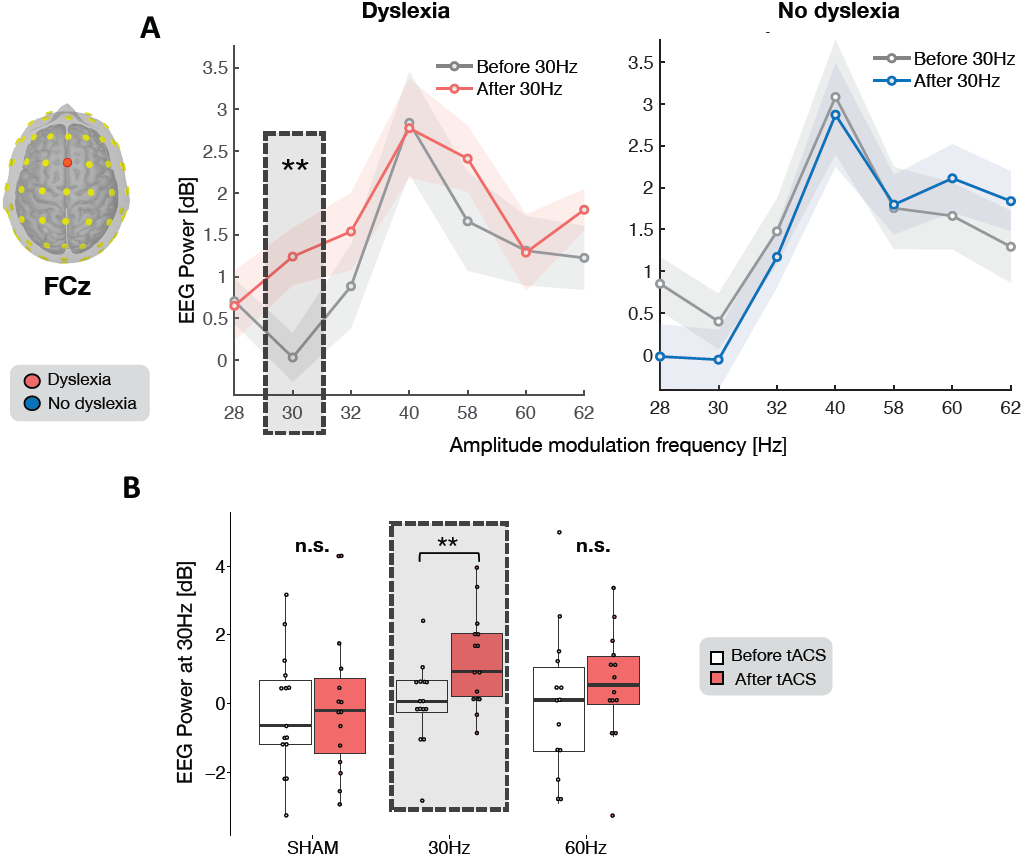
tACS-induced power modulation in auditory steady-state response at the scalp level. Auditory steady-state response in power (dB) to pure-sounds modulated in amplitude (AM) with specific frequencies (from 28 Hz to 62 Hz, x-axis) recorded at electrode FCz. This electrode was chosen as it displayed the strongest evoked response in the time domain (Fig. **1**). For each amplitude modulation frequency, we considered the power response at the specific frequency used to modulate the pure tones stimuli. Both the dyslexia (**A**, left) and no-dyslexia group (**A**, right) showed the maximal response evoked by the auditory stimulus modulated with a 40 Hz envelope, and the minimum power elicited by the 30 Hz AM tones. In the dyslexia group, the 30 Hz-tACS elicited a selective power increase for the 30 Hz AM sounds at 30 Hz (**A**, * for p < 0.05, ** for p < 0.01). This effect was absent in the sham and in the 60 Hz condition in the same group (**B**), and in the no-dyslexia group (**A**, right).

Having established that subjects with dyslexia had a selective deficit in neural oscillations at 30 Hz, we examined the impact of tACS on behavior. We first assessed the effect of each stimulation condition on phonemic awareness, considering the number of errors in the repeated phonemes at the pseudo-word test. We found that, unlike either control conditions (60 Hz and sham), 30 Hz tACS elicited a significant interaction between group (dyslexia/no dyslexia) and session (before/after/one hour after, F_2,56_ = 10.56, p < 0.001, η^2^_p_ = 0.27, Fig **2A**, middle panel). Following up on this result, we analysed performance separately in the two groups and found a main session effect only in the dyslexia group (dyslexia: F_2,28_ = 7.51, p < 0.01, η^2^_p_ = 0.35; no dyslexia: F_2,28_ = 3.31, p > 0.05, η^2^_p_ = 0.19). Post-hoc t-tests revealed that phonemic awareness significantly improved *after* 30-Hz stimulation (T_14_ = 3.78, p < 0.01, FDR corrected, d = 0.98, Fig **2A**), but decreased back to baseline *one hour after* the stimulation (T_14_ = -2.56, p < 0.05, FDR corrected, d = 0.66, Fig **2A**). As expected, adults without dyslexia performed better than those with dyslexia across all tACS conditions (*sham*: F_1,28_ = 24.42, p < 0.001, η^2^_p_ = 0.46; *30 Hz*: F_1,28_ = 17.9, p < 0.001, η^2^_p_ = 0.39; *60 Hz*: F_1,28_ = 18.97, p < 0.001, η^2^_p_ = 0.4, Fig **2A**).

Although several models agree on a phonological deficit in dyslexia, it is still debated whether the underlying core impairment arises at the phonemic or syllabic level [26]. To address the specificity of the phonemic improvement after 30 Hz stimulation, we also evaluated syllable short-term memory from the pseudo-word test. We compared the number of omitted or wrongly repeated syllables, *before* and *immediately after* the 30 Hz tACS, and found that tACS did not improve syllable processing (T_14_ = 0.43, p > 0.05, d = 0.11, Fig **2B**). Note, however, that since we only probed an effect of 30 Hz tACS, this result does not contradict the hypothesis that slower brain rhythms also play a role in dyslexia by interfering at the syllabic level [37,38].

To further characterize the phonological improvement, we correlated the gain in phonemic awareness induced by 30 Hz tACS with language skills as measured by the ECLA16+. We found a markedly negative relationship (r = -0.48, p < 0.01, Fig **2C**), indicating that the intervention was more effective in participants with more severe dyslexia, even though there was a certain degree of variability, including significant effects in some individuals diagnosed with relatively mild dyslexia.

Finally, to address whether phonemic awareness improvement following 30 Hz-tACS stimulation in the dyslexia group could ultimately result in better reading performance, we extracted and analysed separately reading speed and accuracy from the text reading task. With respect to reading accuracy, we found a main effect of session in the dyslexia group for the 30 Hz-tACS condition (F_2,28_ = 4.35, p < 0.02, η^2^_p_ = 0.23, Fig **2D**), indicating a significant performance improvement after 30-Hz tACS as compared to the session preceding the stimulation (T_14_ = 3.08, p < 0.05, d = 0.8, FDR corrected, Fig **2D**). This effect corresponds to a decrease in error number and dysfluencies, an improvement in accuracy that we also found in the spoonerism test (Fig **S2-A**, left). This effect was not present in normo-readers (F_2,28_ = 1.63, p > 0.05, η^2^_p_ = 0.1). As for the phonemic improvement, the gain in reading accuracy correlated with the ECLA16+ score across groups (reading accuracy: r = -0.50, p < 0.01, Fig **2E**; spoonerism accuracy: r = -0.39, p = 0.03, Fig **S2-C**).

For reading speed, we found a session effect in both groups (dyslexia: F_2,28_ = 5.96, p < 0.01, η^2^_p_ = = 0.3, no dyslexia: F_2,28_ = 7.6, p < 0.01, η^2^_p_ = = 0.35, Fig **S3-A**) indicating that reading speed was slowed-down immediately after all stimulation conditions, regardless of dyslexia severity. Focusing on the 30 Hz-tACS, the non-specificity of the effect was confirmed, as it was not correlated with the ECLA16+ score (r = 0.32, p > 0.05, Fig **S3-B**). Like the facilitating effect on reading accuracy, the reading speed slowdown did not reflect a speed/accuracy trade-off as the two variables were independent (r = 0.01, p > 0.05, Fig **S3-C**). These results suggest that the improvement of phonological awareness directly translated on reading accuracy, while reading speed was altered by a non-specific effect of tACS stimulation. Corroborating this finding, results from the spoonerism test did not show changes in response speed (Fig **S2-B**, left), indicating that 20 min tACS are not sufficient to positively impact speech production speed.

Having established that subjects with dyslexia can benefit from 30 Hz-tACS, we examined the neural activity underpinning the behavioral improvement occurring selectively at this frequency of stimulation.

To take advantage of a maximal S/N we first used electrode FCz, the sensor displaying the strongest evoked response (Fig **1A**), to assess tACS effects on the 30 Hz auditory response separately in the two groups. TACS at 30-Hz elicited a significant power increase in the 30 Hz ASSR in the dyslexia group (T_14_ = 3.7, p < 0.01, d = 0.97, Fig **3A**, left), while no effect was found in controls (T_13_ = 0.8, p = 0.43, d = 0.21, Fig **3A**, right). Critically, we found no enhancement of ASSR power in either group after 60 Hz (T_14_ = -0.991, p > 0.05, d = 0.256) or sham (T_14_ = -0.458, p > 0.05, d = 0.118) tACS (Fig **3B**). Overall, these results indicate that 30 Hz tACS was selectively effective at the frequency that was disrupted in the group with dyslexia, and the absence of such an effect in controls shows that 30 Hz tACS is ineffective if the oscillatory activity is already present.

We further evaluated the effect of 30-Hz tACS on the 30-Hz ASSRs in the dyslexia group separately in each of the two above mentioned ROI (i.e. auditory cortex and superior temporal gyrus) by performing a 2×2 repeated measures ANOVA with session (before/after) and hemisphere (left/right) as factors. In the auditory cortex, we found a significant power increase after 30-Hz tACS (main effect of session, F_1,14_ = 13.71, p < 0.01, η^2^_p_ = 0.49, Fig **4A**). This effect was driven by a significant increase in the left auditory cortex response (T_14_ = 3.4, p < 0.05, d = 0.88, FDR corrected, Fig **4A**), while in the right hemisphere, the difference *before* versus *after* tACS was not significant (T_14_ = 1.6, p > 0.05, d = 0.42, FDR corrected, Fig **4A**). In the STG, the same analysis showed a significant interaction between session and hemisphere (F_1,14_ = 10.6, p < 0.01, η^2^_p_ = 0.43, Fig **4B**): like in the auditory cortex, 30-Hz tACS increased the response in the left hemisphere (T_14_ = 2.8, p < 0.05, d = 0.72, FDR corrected), while the right STG displayed a trend in the opposite direction (T_14_ = -1.5, p = 0.1, d = 0.4, FDR corrected). Interestingly, after 30-Hz tACS, the ASSRs were significantly stronger in the left than the right STG (T_14_ = -3.1, p < 0.05, d = 0.8, FDR corrected), reversing the pattern we observed before stimulation (T_14_ = 1.8, p = 0.12, d = 0.45, FDR corrected). These results indicate that tACS did not only boost 30-Hz neural activity in left auditory regions, but also re-instated its left hemispheric dominance.

**Fig. 4.**
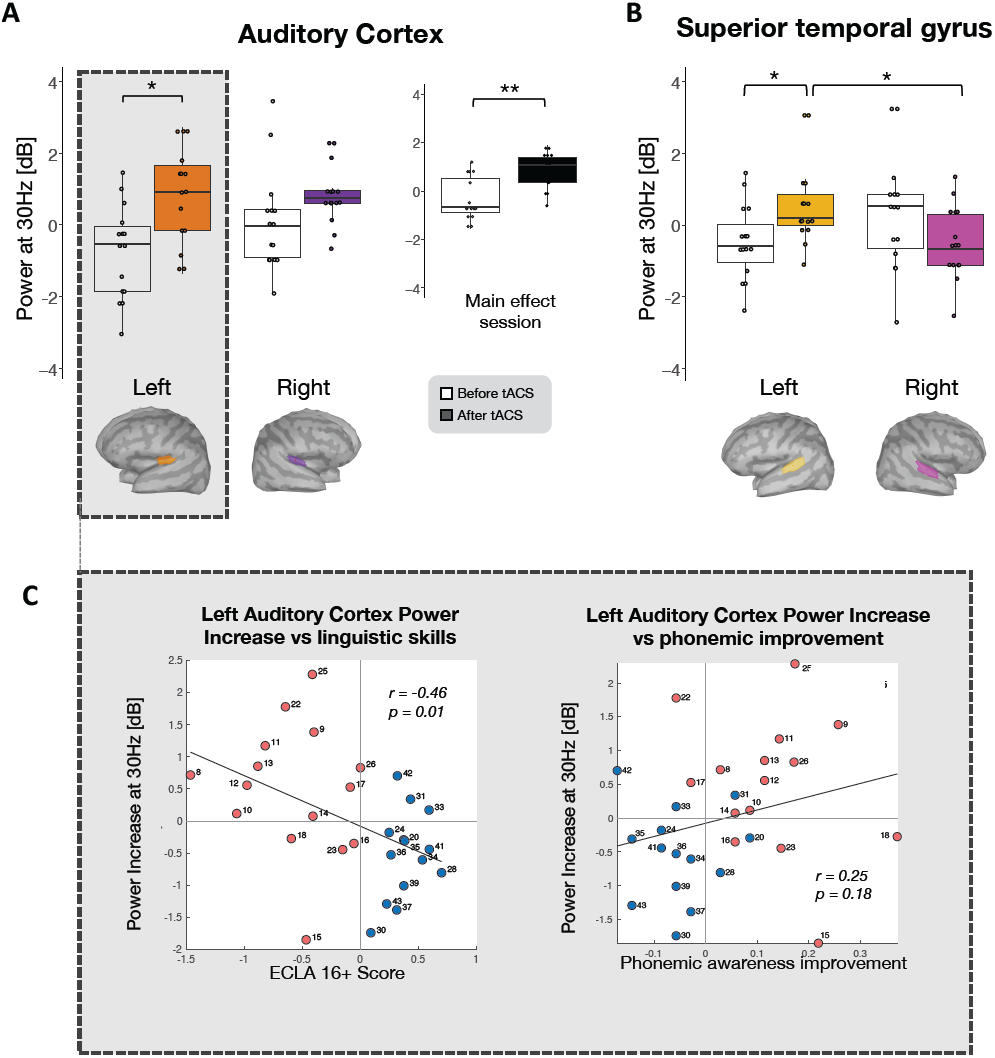
Power increase after 30-Hz tACS in the dyslexia group selective to the left hemisphere and relationship with behavioral measurements. Average power (dB) of auditory steady-state responses to 30-Hz amplitude-modulated pure tones over two regions of interest (ROIs) in each hemisphere, *before* (white whisker plots) and *after* (colored whisker plots) 30-Hz tACS. Differences were tested by considering as fixed effects *session* (before/after tACS) and *hemisphere* (left/right) separately in the auditory cortex (**A**) and the superior temporal gyrus (**B**). In auditory cortex, 30-Hz tACS increased responses bilaterally (**A**-right, average activity of both hemispheres for each *session*, main effect of *session*), an effect driven by a significant power increase in the left hemisphere (**A**-left). A significant interaction between *session* and *hemisphere* in the superior temporal gyrus (**B**) revealed that, on the left hemisphere, the response after tACS was significantly higher than the response before tACS as well as the response on the right hemisphere after the stimulation. We considered the change in power at 30 Hz in the left auditory cortex in both the dyslexia and no-dyslexia group and found a negative correlation between the power increase and the severity of dyslexia (**C**-left), while the correlation with the phonemic improvement after 30 Hz-tACS showed a positive trend (**C**-right, * for p < 0.05).

In the control group, the same analyses performed in both ROIs did not reveal a significant difference between hemisphere and session. We then sought for a relationship between the severity of dyslexia and the gain in the auditory cortex and found a negative correlation in the left (r = -0.46, p = 0.013, Fig **4C**, left) but not the right hemisphere (r = 0.14, p = 0.44). We also observed a trend indicating that the phonemic awareness gain was related to the 30 Hz ASSR power increase (r = 0.25, p = 0.18, Fig **4C**, right).

## Discussion

We show that tACS delivered at 30 Hz could enhance phonological abilities in individuals with dyslexia. This finding has first important implications in basic aspects of speech processing research as the relation between low-gamma neural activity and phonemic encoding has so far only been either conjectured [18,39,40] or inferred using correlational studies [22,41,42]. Selectively reinstating low-gamma auditory activity in subjects in whom it is disrupted offers a unique opportunity to address this important issue. By showing that 30 Hz but not 60 Hz tACS can boost phonemic but not syllabic perception, we can now confidently argued for a causal link between the presence of low-gamma activity in left auditory cortex and basic phoneme encoding via temporal integration windows of about 35ms [18].

This results evidently also constitutes an interesting promise for potentially normalizing phonological processing in subjects with dyslexia. Previous attempts at improving reading efficiency and phonological processing using neuromodulation already gave encouraging results [6–8]. However, as these studies used transcranial direct current stimulation (tDCS) or other neuromodulation techniques that do not target specific brain rhythms and neural processes the exact neural bases of performance enhancement could not be specified. The current results show that phonological processing improved via a focal and specific enhancement of 30-Hz oscillatory activity in left auditory cortex. This finding confirm others showing that tACS can indeed influence neuronal activity in a frequency- and location-specific manner [43,44]. Phonological performance improvement hence likely reflects changes in neuronal entrainment towards the frequency of the tACS stimulation [43,45].

Interestingly, the facilitating tACS effect on phonological processing was only observed in subjects with dyslexia, and was proportional to dyslexia severity as indexed by the ECLA score. This observation also confirms previously observed performance-dependent effects of transcranial electrical stimulation, manifesting as a larger behavioral improvement in poorly performing relative to proficient individuals [46–48]. As observed experimentally [49] and predicted by computational models [31,50], tACS efficacy depends on the gain in endogenous oscillation power. The variable effect depending on initial phonological performance reported here could hence denote the initial difference in 30 Hz power gain in left auditory cortex between the two groups [22], as confirmed by the current data. From a neurobiological perspective, tACS performance-dependent effectiveness might reflect individual changes in the excitatory/inhibitory balance, a variable that has previously associated with reading performance [51]. The presence of variable levels of dysregulation of cortical excitability in dyslexia could explain that the same stimulation did not lead to comparable effects in all subjects, and that there was no significant oscillation power gain in the control group.

Reinstating 30 Hz activity had an immediate impact on the phonological deficit, as it improves phonemic processing (pseudoword reading, spoonerism, phonemic accuracy in text reading) by about 10 to 15%, However, a single 20 minutes tACS exposure was not sufficient to induce a long lasting low-gamma power boosting effect. Short-lived effects (< one hour) following acute tACS intervention are extensively reported in the literature [52–54]. They are agued to reflect short-term plasticity (STP), which declines over a period of half an hour [55,56]. STP selectively amplifies synaptic signals in a frequency-dependent manner, presumably inducing spike-timing dependent plasticity via N-methyl-D-aspartate (NMDA) receptors [55–57]. However, whether the beneficial tACS effects on phonological processing globally reflects a normalization of glutamatergic activity, previously shown to be excessive in poor readers [51], remains to be demonstrated.

Short-term plasticity *per se* is not sufficient to induce long-term potentiation (LTP) lasting from hours to days [55,56]. Longer lasting changes can be obtained by repeating the tACS simulation over several consecutive days [58]. With tDCS, cumulative effects can be observed after only three daily 20 min exposure [59], and can last up to one months after the first session. A 6 months effect has been observed after 6 days of tACS combined with targeted behavioral training [60]. We could hence expect to induce a longer-lasting improvement of phonemic processing, and to stabilize tACS effects through consecutive, repetitive stimulation sessions, and these effects could even be potentiated by combining tACS administration with a phonological training [61].

Applied in children, such a chronic treatment might holds the potential to not only durably normalize the phonemic sampling capacity, but also induce large scale modifications of the reading network [12].

Our results already suggest that larger-scale effect could be induced, as tACS also induced a re-lateralization of 30-Hz responses to the left temporal cortex. A similar immediate interhemispheric effect was previously reported in the context of interactions between primary motor cortices using other neuromodulation techniques [62,63]. If stabilized by repeated stimulation, this large-scale effect might also confer the 30 Hz tACS procedure the interesting advantage of (re)circumscribing phonemic processing to the left hemisphere, an important functional property for improving reading speed in the longer run. Future protocols might also exploit dose-dependent neural entrainment by tACS to tailor stimulation intensity to individual neural deficit in gamma activity [44].

An important finding of the current study is that tACS only improved reading accuracy but not reading speed. This discrepancy did not appear to be due to a speed/accuracy trade-off, and more likely reflect that the acute 20 min. stimulation acted primarily on phonemic encoding locally in bilateral temporal cortices, but not on phonological downstream processing. The access to phonemic representation via interactions with the left inferior frontal cortex remained presumably unchanged by the stimulation. Such critical interactions could however improve if the phonemic representation format were durably normalized by repeated tACS: if this is the case, an improvement of reading speed could be expected, in accordance with the phonological deficit hypothesis.

The current results demonstrate for the first time the causal role of low-gamma oscillatory activity in phonemic processing, and further show the selective impact of targeted tACS on the phonological deficit in dyslexia. This new line of research offers interesting perspectives for promoting plasticity in the reading network, via the correction of basic properties of the auditory cortex [64].

## Methods

### Participants

Fifteen adults previously diagnosed with dyslexia (*dyslexia group*, 13 women, 2 men, mean age 27.4 years, SD ± 9, range 18-47) and fifteen fluent readers (*no-dyslexia group*, 11 women, 4 men, mean age 25.6 years, SD ± 7.8, range18-47) took part in our study (Table **S2** and Fig **S5**).

The experimental paradigm was approved by the local Ethics Committee (*Commission cantonale d’éthique de la recherche*, project #15-264), and was performed in accordance with the Declaration of Helsinki. The study has been registered retrospectively in a publicly accessible clinical trials registry approved by the WHO on www.clinicaltrials.gov (NCT04277351). All participants provided written informed consent and received a financial compensation for their participation.

The exclusion criteria were a history of brain injury, neurological or psychiatric disorders, and the presence of invasive electronic implants. All participants had normal hearing acuity as assessed with an audiogram (pure-tone frequency threshold between 250 and 8000 Hz) and adequate nonverbal intelligence (standard score above 80 on Raven’s progressive matrices, Raven, Court, & Raven, 1984). The two groups did not present any difference in non-verbal intelligence (T_28_ = 1.37, p > 0.05, d = -0.5). To be included in the *dyslexia group*, participants were required to present a history of dyslexia previously assessed by a speech-language therapist, that had to be confirmed by the ECLA16+ test during the *inclusion day*.

### Experimental paradigm

The protocol was spread over four experimental days (Fig **S1**): an *inclusion day* to assess language and cognitive performance and three *experimental days* during which transcranial alternating current stimulation (tACS) was administered. The tests lasted approximately 5-6h/day, amounting to a total duration of 22 hours of experimental time per participant (Fig **S1**).

The auditory stimuli used in the language tests and during the EEG experiment were delivered binaurally using insert earphones (ER-2, Etymotic® Research Inc., Elk Grove Village, IL) at 70–75 dB SPL via a graphical user interface developed in Matlab (version 2015). Responses to the language tests were recorded with a microphone and subsequently analyzed by a certified linguist.

#### Inclusion day (day 1)

The dyslexia diagnosis was confirmed during the *inclusion day* using the ECLA16+ test, a standardized tool to evaluate reading proficiency (positive values indicates high performance). The test includes multiple sub-tests, assessing phonological awareness, short term memory and reading skills. Further statistical analysis (two-tailed unpaired t-test between groups) confirmed that the dyslexia group performed worse than fluent readers for all individual skills tested (phonological awareness: T_28_ = 7.6, p < 0.0001, verbal short term memory: T_28_ = 4.2, p < 0.001, reading skills: T_28_ = 4,9, p < 0.0001, Table **S2**). Subjects additionally performed a rapid automatized naming (RAN) test, which confirmed that the dyslexia group had reduced lexical access (two-tailed unpaired t-test between groups, T_28_ = 5.62, p < 0.0001, Table **S2**). Last, for familiarization purposes, participants underwent a training session with the same battery of custom-designed language tests as those used during the experimental days (see below).

#### Experimental days (days 2-4)

During each of the following *experimental days (*day 2 to 4), participants received one of three tACS stimulation conditions (30 Hz, 60 Hz and a sham condition where subjects received no stimulation), in a counterbalanced order across subjects. Experimental days were separated by at least ten days. Each *experimental day* included three experimental sessions, respectively *before, after*, and *1h after* tACS, in order to reveal both immediate and potentially longer-lasting tACS effects. At the beginning of each of the three sessions, we recorded auditory steady-state responses (ASSR) to pure tones modulated in amplitude at various frequencies (28-62 Hz) by means of electroencephalography, followed by language tests.

### Language assessment

Language tests were specifically designed for this study by a certified linguist in order to probe those skills that are most strongly impaired in dyslexia, namely phonological awareness, short-term memory, as well as reading speed and fluency. These skills were assessed using three tests: pseudo-word repetition, spoonerism and text reading (Table **S1**). The development of a custom-designed solution was necessary to ensure the diversity of linguistic material across experimental sessions (*before, after, 1h after*) and minimise learning effects that may occur over repeated sessions and experimental days. All versions of each language test were matched for difficulty, i.e. similar phonological and syntactic features, same lexical frequency across sessions. To evaluate whether a remaining learning effect could occur over the three experimental sessions of one experimental day despite these precautions, we recruited twenty-five fluent-readers (not included in the main experiment) and had them perform the three sessions of language tests in the same order and with the same timing as the ones used in the experimental paradigm. A one-way repeated-measures analysis of variance (ANOVA) did not reveal any statistically significant effect of session for any of the three tests.

#### Pseudo-word repetition

This test assesses the participant’s phoneme representation (i.e. the ability to retrieve and recall each phoneme) and syllabic short-term memory (i.e. the ability to recall syllable sequences as accurately as possible, Pickering, 2006; Snowling, 2013). A total of 30 different items were included in the test; participants were instructed to repeat single pseudo-words immediately after hearing them. We evaluated *phonemic awareness* by considering errors in phonemes repetition due to a perception deficit, that is regarding a single articulatory feature (voice onset timing or place of articulation). *Syllable short-term memory* was computed by taking into account the number of non- or wrongly repeated syllables.

#### Spoonerism

This task assesses phonological and lexical structure awareness [67,68]. Participants were presented with two regular words and were asked to repeat them after transposing the first phoneme (e.g. hand – pig, becomes pand – hig). From this test we calculated an accuracy index as the total number of correctly inverted words, and a speed score as the average time required to perform one phoneme inversion. A *global phonological awareness* score was computed by averaging the two above mentioned indexes.

#### Text reading

This test evaluates reading fluency, lexical knowledge (written word identification) and the ability to convert lexical orthography to phonology [69]. Participants were asked to read as fluently as possible a scientific text for three minutes [70]. *Reading speed* corresponded to the number of read words and *reading fluency* to the number of errors and dysfluencies.

### EEG recording and stimuli

Along with the language tests, we recorded EEG using a 64-channel setup (Brain Products GmbH, Gilching, Germany) *before, immediately after*, and *1h after* each tACS condition (30 Hz, 60 Hz, and sham). As the EEG cap was not removed between sessions, a passive-electrode EEG system was chosen to ensure stable impedances throughout the entire experimental day (∼6h). Electrode AFz was used as ground contact and FCz as reference. Raw signals were sampled at 1 kHz using proprietary software (Recorder, BrainProducts GmbH).

During the EEG recordings, we presented amplitude-modulated (AM) sounds to entrain brain oscillations in a frequency-specific manner (frequency-tagging) and measured the resultant ASSR in the steady period beginning 500 ms after sound onset and following the initial auditory-evoked potential. Frequency-tagging probes the capacity of auditory cortical regions to synchronize to an auditory stimulus with constant amplitude [71,72]. Pure tones (carrier frequency: 1000 Hz) modulated in amplitude at specific frequencies (28, 30, 32, 40, 58, 60, 62 Hz) were presented for 1.5 seconds. For each AM condition, the stimulus presentation was repeated 40 times with a 3.5-second inter-stimulus interval. Each EEG-ASSR block lasted approximately 25 minutes, during which participants remained seated in front of a screen placed 1 meter away from their forehead, displaying a muted video of their choice. In order to minimize artifacts on the EEG traces, participants were asked to avoid eye and body movements.

### Transcranial alternating current stimulation

Transcranial alternating current stimulation (*Starstim*, Neuroelectrics, Spain) was delivered via five electrodes (*Pistim*, Neuroelectrics, Spain, 1.2 cm diameter) placed over the left auditory cortex and integrated into the EEG cap to ensure invariant position between participants. Electroconductive gel was used to ensure optimal conductance between the electrodes and the scalp. Electrodes were arranged in a 4×1 ring configuration, at TTP7h, FTT9h, FCC5h, CPP5h, TPP9h, with the central anode delivering an alternating current below 2 mA, and the surrounding four cathodes delivering ¼ of the anode’s current in the opposite polarity (Fig **S1**). The 4×1 electrode configuration is a well-established experimental procedure that delivers the electrical stimulation focally to a specific brain region [73].

The intensity of tACS stimulation was tuned separately for each participant, starting from 0.8 mA and increasing by steps of 100 µA until perception threshold, with a maximum intensity of 2 mA. The current was then reduced below that threshold and this value was kept constant throughout the entire duration of the experiment, with 20 seconds of ramp-up and down. The tACS stimulation lasted 20 minutes, a common duration in cognitive neuroscience research [32]. In the Sham condition, 30-Hz tACS was delivered only during the ramp-up and down periods (20 seconds); no current was delivered during the 20 minutes intervention.

Overall, subjects’ reports indicated that they could not tell whether they received real or sham tACS: out of 30 participants, only 7 correctly reported having received a placebo stimulation during the sham condition, while 18 participants denied having received any sham tACS and believed that all three experimental days featured active electrical stimulation.

At the end of the stimulation, subjects were debriefed about side effects (pain, tingling and any skin sensation, warming, fatigue or attentional difficulties), associated with tACS by rating the level of discomfort on a scale between 0 and 10 (Fig **S5**). A repeated measures ANOVA with group (dyslexia/no dyslexia) as between-subjects factor and tACS condition (sham/30 Hz/60 Hz) as within-subjects factor revealed a main effect of stimulation condition (F_2,28_ = 9.1, p < 0.001, η^2^_p_ = 0.23). Post-hoc analyses showed weaker reported side effects for the sham than the 30 Hz (T_29_ = 3, p < 0.01, d = 0.54) and 60 Hz conditions (T_29_= 4.3, p < 0.001, d = 0.78). We found no difference in the reported negative perceptions between the two active stimulation groups. Given the absence of significant difference in the reported side effects between 30 Hz and 60 Hz condition, and between groups, we can exclude that the enhanced behavioral performance following the 30 Hz tACS, selective in the dyslexia group, could have been influenced by discomfort sensations.

### Data analysis

#### Statistical analysis of linguistic tests

All behavioural data were rescaled between 0 (low performance) and 1 (high performance) by taking into consideration the entire pool of 30 subjects, separately for each subtest. Even though no learning effect over the three sessions (i.e. *before, after* and *1h after*) within the same day was observed in the pilot behavioural study run in 28 independent subjects, we identified a trend for improvement over experimental days. For this reason, we considered, for each participant and each test separately, the performance during the first session (*before* stimulation) of each experimental day. We computed a slope value reflecting the theoretical improvement that might occur between the first and last experimental day and regressed it out from the dataset.

In order to investigate the impact of tACS over phonemic representation, we analysed the phonemic awareness score separately for each tACS condition by applying a 2×3 repeated measures ANOVA with group (dyslexia/no dyslexia) as between-subjects factor and session (before/ after/1h after) as within-subjects factor. In case of significant effect, we assessed whether changes in phonemic awareness would be also accompanied by variations in syllable processing, and analysed the syllable short term memory for the specific session/tACS condition of interest.

We then analysed indexes of accuracy and speed from the text reading and spoonerism tests. Separately in each group, we applied a 3×3 repeated measures ANOVA with session (before/after/1h after tACS) and stimulation condition (30 Hz/60 Hz/sham) as factors.

We performed pairwise post-hoc comparisons using paired t-tests, correcting for multiple comparisons with the False Discovery Rate (FDR) method. In accordance with our hypothesis, we restricted the comparisons to performance differences between sessions within the same tACS condition.

To investigate whether potential behavioural changes between the sessions before and after tACS stimulation were related dyslexia severity, we performed Pearson correlation analyses between the ECLA 16+ score and the performance difference between the two sessions of interest. We investigated the existence of potential relationship between improvement in reading speed and accuracy between two sessions (trade-off hypothesis), by performing a Person correlation analysis between these two measurements.

#### EEG pre-processing and analysis

EEG data pre-processing was conducted using the EEGlab v14.1.2 [74] and SASICA [75] toolboxes within the Matlab environment. Signals were down-sampled at 500 Hz and filtered using a Hamming windowed sinc FIR filter between 1 and 70 Hz. EEG epochs were defined from 1s pre-stimulus to 2s post-stimulus. Epochs contaminated by strong muscular artifacts were excluded by visual inspection. Noisy channels were automatically identified using a custom-written Matlab script based on the presence of high-frequency activity, inspected visually and subsequently removed. When a single channel exhibited an epoch-specific artefact, the signal was interpolated for that epoch only. Data were then re-referenced to average reference. Independent component analysis (ICA) was applied to the epoched dataset, whose dimensionality was previously reduced by principal component analysis to 32 components. Artifactual components were identified using a semi-automatic method based on, among others, measures of focal channel topography, autocorrelation and generic discontinuity available through the SASICA toolbox. After ICA-based de-noising, the artifact-contaminated channels initially removed were interpolated using spherical splines, all epochs were visually inspected again and rejected if artefacts remained. Data from one participant in the no-dyslexia group was excluded from the analysis due to a strong artefact contamination.

Subsequent data analysis was performed using Fieldtrip [76] and Brainstorm [77] toolboxes, together with custom scripts.

#### Surface EEG space analysis

First, we computed the grand average signal over time for the dyslexia group to identify the auditory evoked potential and the scalp electrode exhibiting the strongest peak-to-peak amplitude. We restricted the surface space analysis to this electrode, FCz. The time-frequency transform in both surface and source spaces was estimated using a discrete Fourier transform (Hanning taper; 1–70 Hz; 1 Hz steps). At the surface level, we quantified the ASSR for each AM condition as the average power at the AM frequency between 500 ms and 1500 ms after sound onset, normalized with respect to the 1s pre-stimulus baseline. For each AM condition, we tested for differences between the session *before* and *immediately after* the stimulation (two-tailed paired t-test).

#### Source space analysis

The distributed source space, consisting in a 15000-vertex mesh of the cortical surface, was obtained from the segmentation of a template MRI (Colin27, MNI). Using the OpenMEEG [78] implementation in Brainstorm software [77], we generated a 3-layer EEG Boundary Element Method model consisting of the inner skull, outer skull and the scalp surfaces, with corresponding conductivity values of 0.33:0.0125:0.33 S/m respectively. Within this model, the source activity of dipoles distributed over the cortical surface was estimated by a minimum-norm approach with noise normalization (dSPM) and constrained orientation (depth weighting order: 0.5, maximal amount: 10; signal-to-noise ratio: 3; noise covariance regularization: 0.1). Spatial smoothing (FWHM 3mm) was applied for displaying the average source activity (Fig **S4**). We outlined two regions of interest (ROIs) of 79-96 vertices in the primary auditory cortex and the superior temporal gyrus for each hemisphere. The average time-frequency spectrum over each ROI was calculated from source activity with the same method as for the surface analysis. To assess whether we could replicate previous findings [22], we tested for differences between the two groups at baseline in the ASSR at 30 Hz, separately in the left and right primary auditory cortex. Subsequent source space analyses were performed to address putative differences between sessions before and after the 30 Hz-tACS. Differences in the laterality of power response were investigated separately for each of the two ROIs with a 2×2 ANOVA, using session (before/after) and hemisphere (left/right) as factors.

## Acknowledgments

We thank P. Mégevand for helpful discussion about the data analysis and comments on the manuscript; S. Martin for comments on the manuscript; C. Pacoret, Prof. Friedhelm Hummel and the Neurostimulation platform of the Biotech Campus for technical advice.

This study has been supported by the Wyss Center for Bio and Neuro Engineering (WCP-006) and Swiss National Science Foundation (320030B_182855, to A.-L.G.).

## Contributions

A.-L.G. conceived the project, A.-L.G., J.N., S.M. and I.M. designed the experiments, J.N. and S.M. conducted the experiment, S.M., J.N., I.M., L.H.A. and A.-L.G. analyzed the data; A.-L.G., S.M., J.N. and J.D. wrote the manuscript;

## References

1. Norton ES, Beach SD, Gabrieli JDE. Neurobiology of dyslexia. Curr Opin Neurobiol. 2015;30: 73–78. doi:10.1016/j.conb.2014.09.007

2. Bakker DJ. Treatment of developmental dyslexia: A review. Dev Neurorehabil. 2006;9: 3–13. doi:10.1080/13638490500065392

3. Alexander AW, Slinger-Constant A-M. Current Status of Treatments for Dyslexia: J Child Psychol Psychiatry Allied Discip. 2004;19: 744–758. doi:10.1177/08830738040190100401

4. Frey A, François C, Chobert J, Velay J, Habib M, Besson M. Music Training Positively Influences the Preattentive Perception of Voice Onset Time in Children with Dyslexia: A Longitudinal Study. Brain Sci. 2019;9: 91. doi:10.3390/brainsci9040091

5. Loo JHY, Rosen S, Bamiou D-E. Auditory Training Effects on the Listening Skills of Children With Auditory Processing Disorder. Ear Hear. 2016;37: 38–47.

6. Rufener K, Krauel K, Meyer M, Heinze H-J, Zaehle T. Transcranial electrical stimulation improves phoneme processing in developmental dyslexia. Brain Stimul. 2019. doi:10.1016/j.brs.2019.02.007

7. Costanzo F, Rossi S, Varuzza C, Varvara P, Vicari S, Menghini D. Long-lasting improvement following tDCS treatment combined with a training for reading in children and adolescents with dyslexia. Neuropsychologia. 2018; 0–1. doi:10.1016/j.neuropsychologia.2018.03.016

8. Heth I, Lavidor M. Improved reading measures in adults with dyslexia following transcranial direct current stimulation treatment. Neuropsychologia. 2015;70: 107–113. doi:10.1016/j.neuropsychologia.2015.02.022

9. Georgiou GK, Papadopoulos TC, Zarouna E, Parrila R. Are auditory and visual processing deficits related to developmental dyslexia? Dyslexia. 2012;18: 110–129. doi:10.1002/dys.1439

10. Ramus F, Rosen S, Dakin SC, Day BL, Castellote JM, White S, et al. Theories of developmental dyslexia: Insights from a multiple case study of dyslexic adults. Brain. 2003;126: 841–865. doi:10.1093/brain/awg076

11. Swan D, Goswami U. Phonological awareness deficits in developmental dyslexia and the phonological representations hypothesis. J Exp Child Psychol. 1997;66: 18–41. doi:10.1006/jecp.1997.2375

12. Boets B, Wouters J, Van Wieringen A, Ghesquière P. Auditory temporal information processing in preschool children at family risk for dyslexia: Relations with phonological abilities and developing literacy skills. Brain Lang. 2006;97: 64–79. doi:10.1016/j.bandl.2005.07.026

13. Boets B, Op De Beeck H, Vandermosten M, Scott SK, Gillebert CR, Mantini D, et al. in Adults with Dyslexia. 2013; 1251–1255. doi:10.1126/science.1244333

14. Lieder I, Adam V, Frenkel O, Jaffe-Dax S, Sahani M, Ahissar M. Perceptual bias reveals slow-updating in autism and fast-forgetting in dyslexia. Nat Neurosci. 2019;22: 256–264. doi:10.1038/s41593-018-0308-9

15. Vellutino FR, Fletcher JM, Snowling MJ, Scanlon DM. Specific reading disability (dyslexia): What have we learned in the past four decades? J Child Psychol Psychiatry Allied Discip. 2004;45: 2–40. doi:10.1046/j.0021-9630.2003.00305.x

16. Schulte-Körne G, Deimel W, Bartling J, Remschmidt H. The role of phonological awareness, speech perception, and auditory temporal processing for dyslexia. Eur Child Adolesc Psychiatry. 1999;8 Suppl 3: 28–34. doi:10.1007/PL00010690

17. Steinschneider M, Volkov IO, Noh MD, Garell PC, Howard MA. Temporal encoding of the voice onset time phonetic parameter by field potentials recorded directly from human auditory cortex. J Neurophysiol. 1999;82: 2346–57. doi:10.1152/jn.1999.82.5.2346

18. Giraud AL, Poeppel D. Cortical oscillations and speech processing: Emerging computational principles and operations. Nat Neurosci. 2012;15: 511–517. doi:10.1038/nn.3063

19. Joliot M, Ribary U, Llinas R. Human oscillatory brain activity near 40 Hz coexists with cognitive temporal binding. Proc Natl Acad Sci. 1994;91: 11748–11751. doi:10.1073/pnas.91.24.11748

20. Lizarazu M, Lallier M, Molinaro N, Bourguignon M, Paz-alonso PM, Lerma-usabiaga G, et al. Developmental Evaluation of Atypical Auditory Sampling in Dyslexia: Functional and Structural Evidence. 2015;5002: 4986–5002. doi:10.1002/hbm.22986

21. Di Liberto GM, Peter V, Kalashnikova M, Goswami U, Burnham D, Lalor EC. Atypical cortical entrainment to speech in the right hemisphere underpins phonemic deficits in dyslexia. Neuroimage. 2018;175: 70–79. doi:10.1016/j.neuroimage.2018.03.072

22. Lehongre K, Ramus F, Villiermet N, Schwartz D, Giraud AL. Altered low-gamma sampling in auditory cortex accounts for the three main facets of dyslexia. Neuron. 2011;72: 1080–1090. doi:10.1016/j.neuron.2011.11.002

23. Lallier M, Molinaro N, Lizarazu M, Bourguignon M, Carreiras M. Amodal Atypical Neural Oscillatory Activity in Dyslexia. Clin Psychol Sci. 2017;5: 379–401. doi:10.1177/2167702616670119

24. Van Hirtum T, Ghesquière P, Wouters J. Atypical neural processing of rise time by adults with dyslexia. Cortex. 2019;113: 128–140. doi:10.1016/j.cortex.2018.12.006

25. Goswami U. Speech rhythm and language acquisition: an amplitude modulation phase hierarchy perspective. Ann N Y Acad Sci. 2019;1453: 67–78. doi:10.1111/nyas.14137

26. Goswami U. A temporal sampling framework for developmental dyslexia. Trends in Cognitive Sciences. Elsevier Ltd; 2011. pp. 3–10. doi:10.1016/j.tics.2010.10.001

27. Lehongre K, Morillon B, Giraud A-L, Ramus F. Impaired auditory sampling in dyslexia: further evidence from combined fMRI and EEG. Front Hum Neurosci. 2013;7: 1–8. doi:10.3389/fnhum.2013.00454

28. Vanderauwera J, Altarelli I, Vandermosten M, De Vos A, Wouters J, Ghesquière P. Atypical Structural Asymmetry of the Planum Temporale is Related to Family History of Dyslexia. Cereb Cortex. 2016;28: 63–72. doi:10.1093/cercor/bhw348

29. Opitz A, Falchier A, Yan C-G, Yeagle EM, Linn GS, Megevand P, et al. Spatiotemporal structure of intracranial electric fields induced by transcranial electric stimulation in humans and nonhuman primates. Sci Rep. 2016;6: 31236. doi:10.1038/srep31236

30. Herrmann CS, Strüber D, Helfrich RF, Engel AK. EEG oscillations: From correlation to causality. Int J Psychophysiol. 2016;103: 12–21. doi:10.1016/j.ijpsycho.2015.02.003

31. Merlet I, Birot G, Salvador R, Molaee-Ardekani B, Mekonnen A, Soria-Frish A, et al. From Oscillatory Transcranial Current Stimulation to Scalp EEG Changes: A Biophysical and Physiological Modeling Study. PLoS One. 2013;8: 1–12. doi:10.1371/journal.pone.0057330

32. Antal A, Alekseichuk I, Bikson M, Brockmöller J, Brunoni AR, Chen R, et al. Low intensity transcranial electric stimulation: Safety, ethical, legal regulatory and application guidelines. Clin Neurophysiol. 2017;128: 1774–1809. doi:10.1016/j.clinph.2017.06.001

33. Steinschneider M, Liégeois-Chauvel C, Brugge JF. Auditory Evoked Potentials and Their Utility in the Assessment of Complex Sound Processing. The Auditory Cortex. Boston, MA: Springer; 2011. pp. 535–559. doi:10.1007/978-1-4419-0074-6

34. Korczak P, Smart J, Delgado R, Strobel TM, Bradford C. Auditory steady-state responses. J Am Acad Audiol. 2012;23: 146–70. doi:10.3766/jaaa.23.3.3

35. Ross B, Borgmann C, Draganova R, Roberts LE, Pantev C. A high-precision magnetoencephalographic study of human auditory steady-state responses to amplitudemodulated tones. J Acoust Soc Am. 2000;108: 679–691. doi:10.1121/1.429600

36. Ma Y, Koyama MS, Milham MP, Castellanos FX, Quinn BT, Pardoe H, et al. Cortical thickness abnormalities associated with dyslexia, independent of remediation status. NeuroImage Clin. 2015;7: 177–186. doi:10.1016/j.nicl.2014.11.005

37. Goswami U, Thomson J, Richardson U, Stainthorp R, Hughes D, Rosen S, et al. Amplitude envelope onsets and developmental dyslexia: A new hypothesis. Proc Natl Acad Sci U S A. 2002;99: 10911–10916. doi:10.1073/pnas.122368599

38. Goswami U, Gerson D, Astruc L. Amplitude envelope perception, phonology and prosodic sensitivity in children with developmental dyslexia. Read Writ. 2010;23: 995–1019. doi:10.1007/s11145-009-9186-6

39. Ghitza O. Linking speech perception and neurophysiology: Speech decoding guided by cascaded oscillators locked to the input rhythm. Front Psychol. 2011;2: 1–13. doi:10.3389/fpsyg.2011.00130

40. Poeppel D. The analysis of speech in different temporal integration windows: Cerebral lateralization as “asymmetric sampling in time.” Speech Commun. 2003;41: 245–255. doi:10.1016/S0167-6393(02)00107-3

41. Gross J, Hoogenboom N, Thut G, Schyns P, Panzeri S, Belin P, et al. Speech Rhythms and Multiplexed Oscillatory Sensory Coding in the Human Brain. PLoS Biol. 2013;11. doi:10.1371/journal.pbio.1001752

42. Kösem A, van Wassenhove V. Distinct contributions of low- and high-frequency neural oscillations to speech comprehension. Lang Cogn Neurosci. 2017;32: 536–544. doi:10.1080/23273798.2016.1238495

43. Krause MR, Vieira PG, Csorba BA, Pilly PK, Pack CC. Transcranial alternating current stimulation entrains single-neuron activity in the primate brain. Proc Natl Acad Sci. 2019;116: 5747–5755. doi:10.1073/pnas.1815958116

44. Johnson L, Alekseichuk I, Krieg J, Doyle A, Yu Y, Vitek J, et al. Dose-Dependent Effects of Transcranial Alternating Current Stimulation on Spike Timing in Awake Nonhuman Primates. bioRxiv. 2019; 696344. doi:10.1101/696344

45. Vieira PG, Krause MR, Pack CC. tACS entrains neural activity while somatosensory input is blocked. bioRxiv. 2019; 691022. doi:10.1101/691022

46. Sarkar A, Dowker A, Cohen Kadosh R. Cognitive Enhancement or Cognitive Cost: Trait-Specific Outcomes of Brain Stimulation in the Case of Mathematics Anxiety. J Neurosci. 2014;34: 16605–16610. doi:10.1523/jneurosci.3129-14.2014

47. Chang C-F, Muggleton NG, Walsh V, Cheng S -k., Hung DL, Juan C-H, et al. Unleashing Potential: Transcranial Direct Current Stimulation over the Right Posterior Parietal Cortex Improves Change Detection in Low-Performing Individuals. J Neurosci. 2012;32: 10554–10561. doi:10.1523/jneurosci.0362-12.2012

48. Santarnecchi E, Muller T, Rossi S, Sarkar A, Polizzotto NR, Rossi A, et al. Individual differences and specificity of prefrontal gamma frequency-tACS on fluid intelligence capabilities. Cortex. 2015;75: 33–43. doi:10.1016/j.cortex.2015.11.003

49. Neuling T, Rach S, Herrmann CS. Orchestrating neuronal networks: sustained after-effects of transcranial alternating current stimulation depend upon brain states. Front Hum Neurosci. 2013;7: 1–12. doi:10.3389/fnhum.2013.00161

50. Ali MM, Sellers KK, Frohlich F. Transcranial Alternating Current Stimulation Modulates Large-Scale Cortical Network Activity by Network Resonance. J Neurosci. 2013;33: 11262–11275. doi:10.1523/jneurosci.5867-12.2013

51. Pugh KR, Frost SJ, Rothman DL, Hoeft F, Del Tufo SN, Mason GF, et al. Glutamate and Choline Levels Predict Individual Differences in Reading Ability in Emergent Readers. J Neurosci. 2014;34: 4082–4089. doi:10.1523/jneurosci.3907-13.2014

52. Benussi A, Koch G, Cotelli M, Padovani A, Borroni B. Cerebellar transcranial direct current stimulation in patients with ataxia: A double-blind, randomized, sham-controlled study. Mov Disord. 2015;30: 1701–1705. doi:10.1002/mds.26356

53. Lefebvre S, Thonnard JL, Laloux P, Peeters A, Jamart J, Vandermeeren Y. Single session of dual-tdcs transiently improves precision grip and dexterity of the paretic hand after stroke. Neurorehabil Neural Repair. 2014;28: 100–110. doi:10.1177/1545968313478485

54. Kasten FH, Dowsett J, Herrmann CS. Sustained aftereffect of α-tACS lasts up to 70 min after stimulation. Front Hum Neurosci. 2016;10: 1–9. doi:10.3389/fnhum.2016.00245

55. Volianskis A, Collingridge GL, Jensen MS. The roles of STP and LTP in synaptic encoding. PeerJ. 2013;2013: 1–13. doi:10.7717/peerj.3

56. Castro-alamancos MA, Connors BW. Short-term synaptic enhancement and long-term potentiation in neocortex. Neurobiology. 1996;93: 1335–1339.

57. Wischnewski M, Engelhardt M, Salehinejad MA, Schutter DJLG, Kuo MF, Nitsche MA. NMDA Receptor-Mediated Motor Cortex Plasticity After 20 Hz Transcranial Alternating Current Stimulation. Cereb Cortex. 2019;29: 2924–2931. doi:10.1093/cercor/bhy160

58. Gebodh N, Esmaeilpour Z, Adair D, Schestattsky P, Fregni F, Bikson M. Transcranial Direct Current Stimulation Among Technologies for Low-Intensity Transcranial Electrical Stimulation: Classification, History, and Terminology. In: Knotkova H, Nitsche MA, Bikson M, Woods AJ, editors. Practical Guide to Transcranial Direct Current Stimulation: Principles, Procedures and Applications. Cham: Springer International Publishing; 2019. pp. 3–43. doi:10.1007/978-3-319-95948-1_1

59. Soekadar SR, Witkowski M, Birbaumer N, Cohen LG. Enhancing Hebbian Learning to Control Brain Oscillatory Activity. Cereb Cortex. 2014; 1–7. doi:10.1093/cercor/bhu043

60. Cohen Kadosh R, Soskic S, Iuculano T, Kanai R, Walsh V. Modulating neuronal activity produces specific and long-lasting changes in numerical competence. Curr Biol. 2010;20: 2016–2020. doi:10.1016/j.cub.2010.10.007

61. Hamilton RH, Kessler SK, Castillo-Saavedra L, Fregni F, Martin D, Loo C, et al. Methodological Considerations for Transcranial Direct Current Stimulation in Clinical Trials. In: Knotkova H, Nitsche MA, Bikson M, Woods AJ, editors. Practical Guide to Transcranial Direct Current Stimulation: Principles, Procedures and Applications. Cham: Springer International Publishing; 2019. pp. 347–377. doi:10.1007/978-3-319-95948-1_12

62. Suppa A, Ortu E, Zafar N, Deriu F, Paulus W, Berardelli A, et al. Theta burst stimulation induces after-effects on contralateral primary motor cortex excitability in humans. J Physiol. 2008;586: 4489–4500. doi:10.1113/jphysiol.2008.156596

63. Vines BW, Nair DG, Schlaug G. Contralateral and ipsilateral motor effects after transcranial direct current stimulation. Neuroreport. 2006;17: 671–674. doi:10.1097/00001756-200604240-00023

64. Hyafil A, Giraud AL, Fontolan L, Gutkin B. Neural Cross-Frequency Coupling: Connecting Architectures, Mechanisms, and Functions. Trends Neurosci. 2015;38: 725–740. doi:10.1016/j.tins.2015.09.001

65. Pickering SJ. Assessment of Working Memory in Children. Work Mem Educ. 2006; 241–271. doi:10.1016/B978-012554465-8/50011-9

66. Snowling MJ. Early identification and interventions for dyslexia: A contemporary view. J Res Spec Educ Needs. 2013;13: 7–14. doi:10.1111/j.1471-3802.2012.01262.x

67. Frederickson N, Frith U, Reason R. Phonological Assessment Battery (Manual and Test Materials). Windsor: N. 1997.

68. Gola-Asmussen C, Lequette C, Pouget G, Rouyer C, Zorman M. ECLA-16+: Evaluation des Compétences de Lecture chez l’Adulte de plus de 16 Ans. 2010.

69. Sprenger-Charolles L, Siegel LS, Béchennec D, Serniclaes W. Development of phonological and orthographic processing in reading aloud, in silent reading, and in spelling: A four-year longitudinal study. J Exp Child Psychol. 2003;84: 194–217. doi:10.1016/S0022-0965(03)00024-9

70. Masmoudi S, Naceur A. Du percept à la décision. Intégration de la cognition, l’émotion et la motivation. De Boeck. 2010.

71. Miyazaki T, Thompson J, Fujioka T, Ross B. Sound envelope encoding in the auditory cortex revealed by neuromagnetic responses in the theta to gamma frequency bands. Brain Res. 2013;1506: 64–75. doi:10.1016/j.brainres.2013.01.047

72. Ross B, Herdman AT, Pantev C. Stimulus Induced Desynchronization of Human Auditory 40-Hz Steady-State Responses. 2005; 4082–4093. doi:10.1152/jn.00469.2005.

73. Datta A, Bansal V, Diaz J, Patel J, Reato D, Bikson M. Gyri-precise head model of transcranial direct current stimulation: Improved spatial focality using a ring electrode versus conventional rectangular pad. Brain Stimul. 2009;2: 201-207.e1. doi:10.1016/j.brs.2009.03.005

74. Delorme A, Makeig S. EEGLAB: An open source toolbox for analysis of single-trial EEG dynamics including independent component analysis. J Neurosci Methods. 2004;134: 9–21. doi:10.1016/j.jneumeth.2003.10.009

75. Chaumon M, Bishop DVM, Busch N a. A Practical Guide to the Selection of Independent Components of the Electroencephalogram for Artifact Correction. J Neurosci Methods. 2015. doi:10.1016/j.jneumeth.2015.02.025

76. Oostenveld R, Fries P, Maris E, Schoffelen JM. FieldTrip: Open source software for advanced analysis of MEG, EEG, and invasive electrophysiological data. Comput Intell Neurosci. 2011;2011. doi:10.1155/2011/156869

77. Tadel F, Baillet S, Mosher JC, Pantazis D, Leahy RM. Brainstorm: A user-friendly application for MEG/EEG analysis. Comput Intell Neurosci. 2011;2011. doi:10.1155/2011/879716

78. Alexandre Gramfort, Théodore Papadopoulo, Emmanuel Olivi, Maureen Clerc. OpenMEEG: opensource software for quasistatic bioelectromagnetics. Biomed Eng Online. 2009;8: 1. doi:10.1186/1475-925X-8-1

