## Supplementary information for "Selective enhancement of low-gamma activity by tACS improves phonemic processing and reading accuracy in dyslexia"

### Supplementary figures

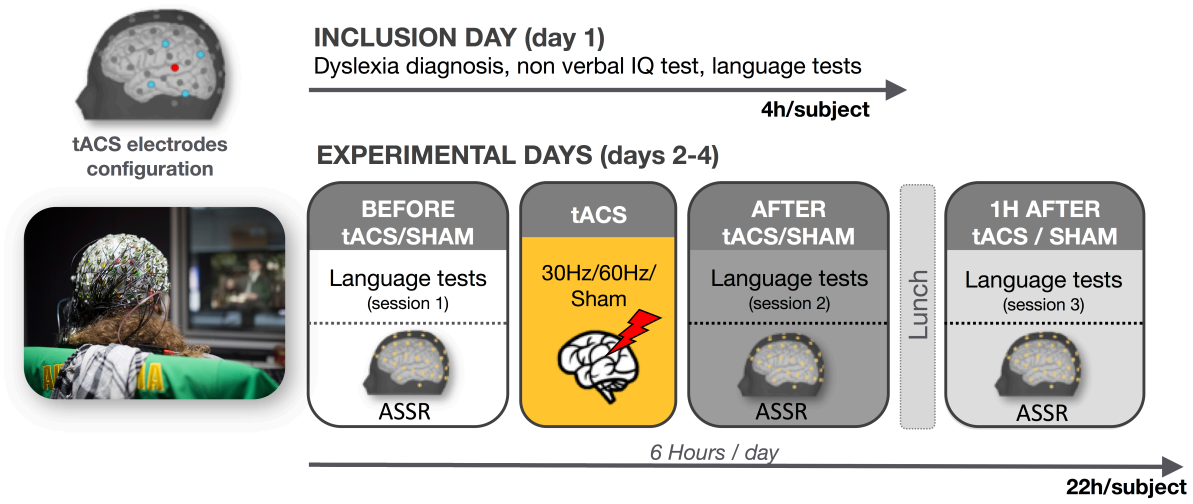


**Fig. S1. Experimental paradigm**. Each participant (15 with dyslexic, 15 non-dyslexics) underwent four recording sessions throughout four experimental days. During the *inclusion day* (day 1) the severity of dyslexia was assessed with ECLA 16+ test together with evaluating baseline performance at three linguistic tests. These included the *pseudo-word repetition test*, *spoonerism test*, and *text reading,* probing among others, errors at the phonemic and syllabic level*.* This allowed us to evaluate phonological awareness, verbal short-term memory, as well as reading fluency, and the ability to convert lexical orthography to phonology. During each of the following experimental days (days 2-4), one of the three tACS stimulation conditions (*sham, 30 Hz and 60 Hz*) was administered, with the order of the tACS condition counterbalanced across subjects. The stimulation lasted 20 minutes and was delivered by means of five tACS electrodes organised as a 4x1 ring and centred over the left auditory cortex. Within each experimental session, performance at the three linguistic tests were evaluated *before, immediately after,* and *one hour after* the tACS stimulation. During these sessions, and for each tACS condition, EEG data were recorded by means of a 64 channels cap. We used auditory steady state EEG responses (ASSR) to amplitude modulated pure-tone sounds with a fixed frequency (from 28Hz to 60 Hz), to entrain brain oscillations in a frequency specific manner.

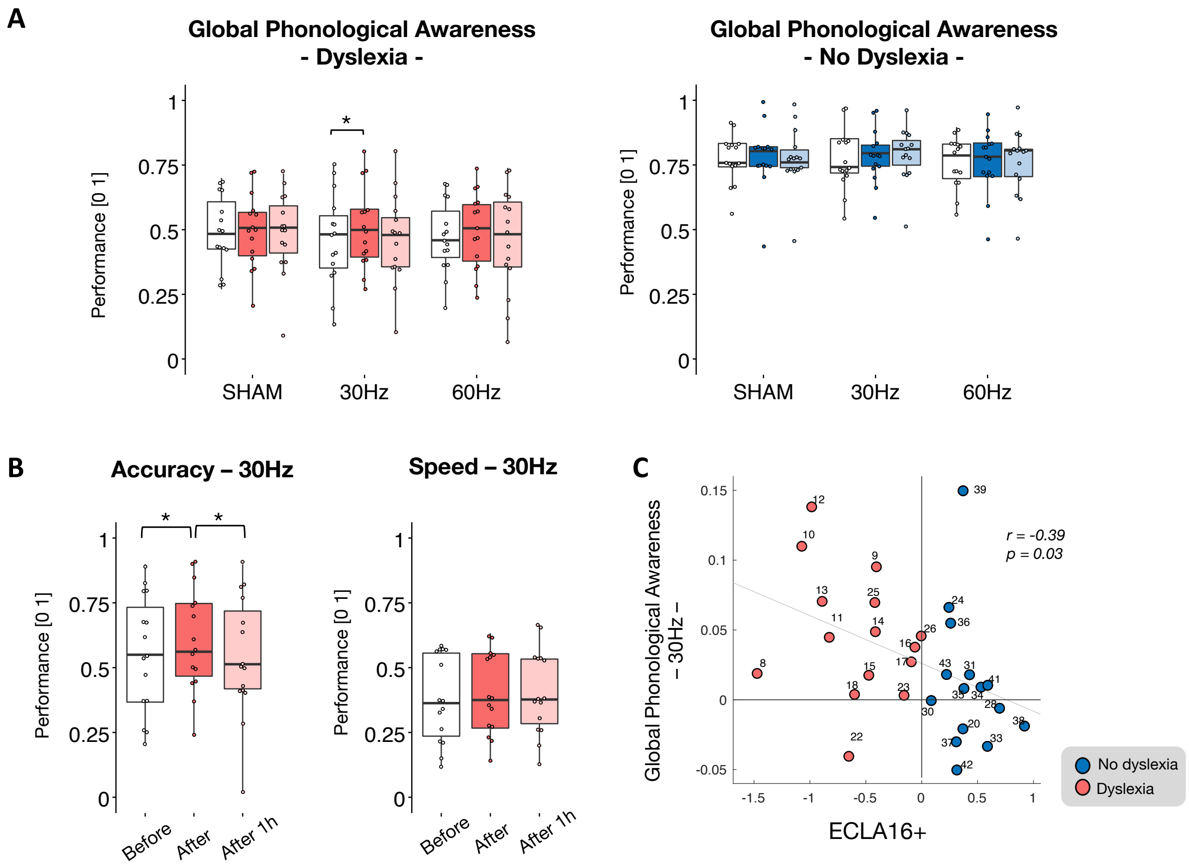


**Fig. S2. tACS stimulation effects on spoonerism test.** We evaluated the global phonological awareness by considering the average of response speed and accuracy at the spoonerism test (**A**). The 3x3 ANOVA with tACS condition (sham/30 Hz/60 Hz) and session (*before*, *after* and *one hour after*) did not revealed any statistically significant effect for neither the dyslexia (**A**, left), nor the no-dyslexia group (**A**, right). However, exploratory post-hoc analysis revealed a performance increase immediately after the stimulation ($T_{14} = 3.86$, $p < 0.05$, FDR corrected, d = 0.99, **A**, left) in the dyslexia group, similarly to the effect we found for reading accuracy and phonemic awareness after the 30 Hz stimulation in the same group. The improvement was stronger in participants with more severe dyslexia (r = -0.39, p = 0.03, **C**). In the dyslexia group, this effect was specific to the response accuracy (mean effect of *session*, $F_{2,28} = 3.95$, $p < 0.05$, $\eta^2_p =  = 0.22$, **B**-left), whereas no difference was present in response speed ($F_{2,28} = 3.28$, $p > 0.05$, $\eta^2_p =  = 0.19$, **B**-right). In particular, response accuracy increased immediately after the 30 Hz stimulation ($T_{14} = 2.63$, $p < 0.05$, d = 0.68) and decreased one hour after the stimulation ($T_{14} = -2.26$, $p < 0.05$, d = 0.58, **B**-left).

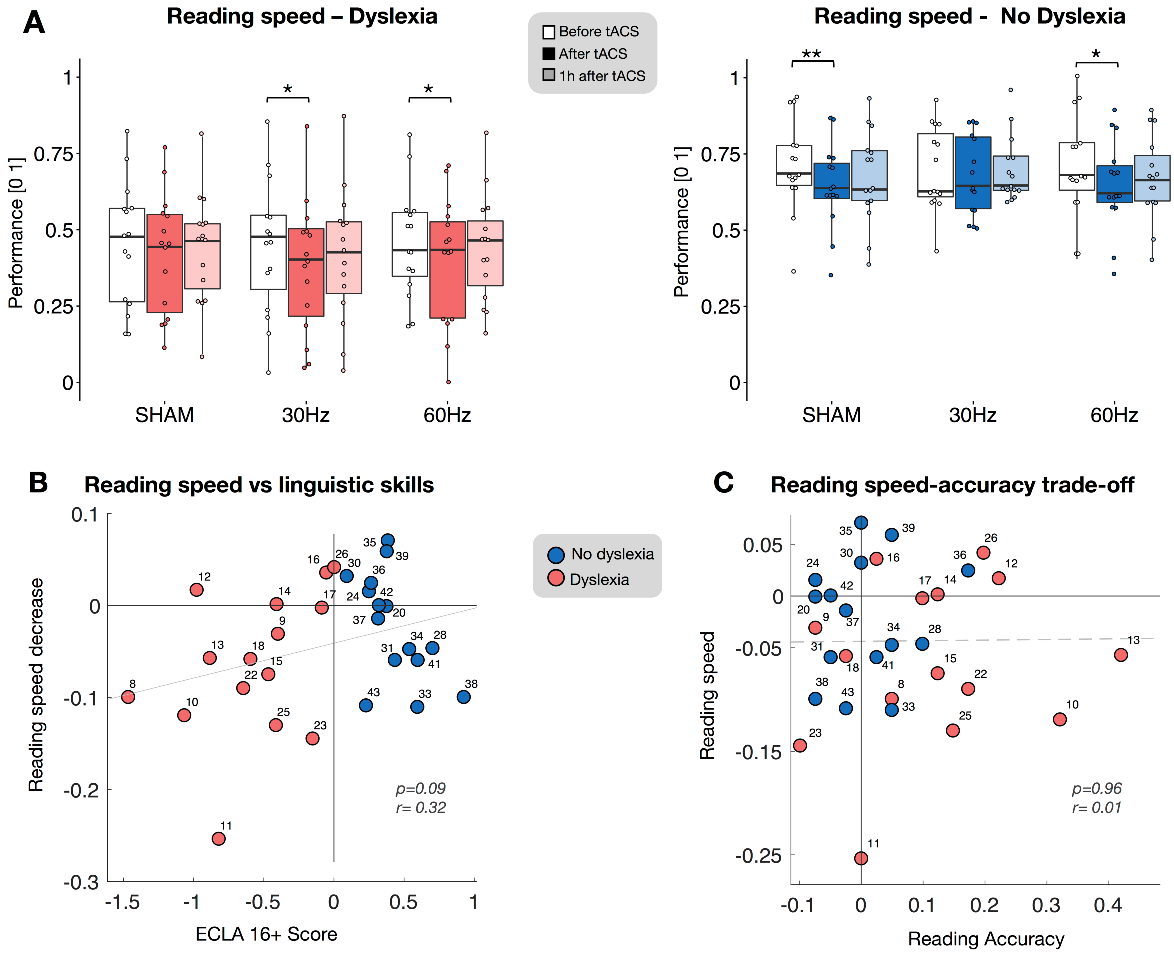


**Fig. S3. tACS stimulation effects on reading speed and speed-accuracy trade-off.** Reading speed performance for each tACS condition (sham, 30 Hz and 60 Hz) and for the sessions *before* (white bar plot)*, after (*dark color*),* and *one hour after tACS* (light color) in the dyslexia (**A**-left) and no dyslexia groups (**A**-right). The ANOVA revealed a main effect of session in both groups (dyslexia: $F_{2,28} = 5.96$, $p < 0.01$, $\eta^2_p =\ = 0.3$, no dyslexia: $F_{2,28} = 7.6$, $p < 0.01$, $\eta^2_p =\ = 0.35$) in the direction of a slowing down immediately after all the stimulation conditions. More specifically, the dyslexia group showed a speed decrease after the 30 Hz stimulation ($T_{14} = -3.1$, $p < 0.05$, FDR corrected, d = 0.8) and 60 Hz tACS ($T_{14} = -3.3$, $p < 0.05$, FDR corrected, d = 0.87). In the no-dyslexia group, reading speed decreased after the SHAM stimulation ($T_{14} = -4$, $p < 0.01$, FDR corrected, d = 1.04) and after 60 Hz-tACS ($T_{14} = -2.9$, $p < 0.05$, FDR corrected, d = 0.76). The correlation between reading speed and ECLA 16+ score did not reveal a significant effect (r = 0.32, p = 0.09, **B**). We assessed a potential speed-accuracy trade-off by measuring the relationship between reading speed and accuracy and we could not find any statistically significant relationship between these two variables (r=0.01, p > 0.05, **C**).

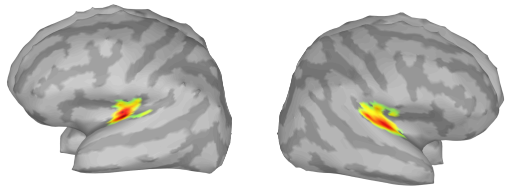

**Fig. S4. Auditory evoked response in the source space**. Source localization revealed that the neural generators of the evoked response at short latencies (~60 ms post-stimulus onset) are located over the primary auditory cortex bilaterally.

| Group | Participant Number | Gender | Age | ECLA16+ | Raven matrix | tACS Side Effects | | |
|---|---|---|---|---|---|---|---|---|
| | | | | | | SHAM | 30Hz | 60Hz |
| Dyslexia | 1 | F | 30 | -1.47 | 108 | 0.83 | 2.20 | 2.17 |
| | 2 | F | 22 | -0.40 | 81 | 6.17 | 4.33 | 4.33 |
| | 3 | F | 19 | -1.07 | 100 | 1.67 | 3.50 | 4.00 |
| | 4 | F | 22 | -0.82 | 102 | 1.67 | 3.33 | 0.67 |
| | 5 | F | 19 | -0.98 | 122 | 2.67 | 2.17 | 4.17 |
| | 6 | F | 20 | -0.88 | 80 | 3.33 | 4.83 | 5.67 |
| | 7 | F | 22 | -0.41 | 115 | 0.83 | 2.50 | 1.67 |
| | 8 | F | 46 | -0.47 | 102 | 3.33 | 1.50 | 4.00 |
| | 9 | M | 30 | -0.06 | 142 | 1.17 | 1.40 | 5.17 |
| | 10 | F | 30 | -0.09 | 81 | 1.00 | 1.67 | 2.33 |
| | 11 | F | 34 | -0.60 | 104 | 1.67 | 2.50 | 3.17 |
| | 12 | F | 24 | -0.65 | 81 | 4.33 | 3.67 | 4.33 |
| | 13 | F | 46 | -0.15 | 142 | 0.83 | 1.50 | 5.17 |
| | 14 | F | 22 | -0.41 | 82 | 1.50 | 2.67 | 3.67 |
| | 15 | F | 24 | 0.00 | 84 | 2.50 | 4.83 | 6.50 |
| No Dyslexia | 16 | F | 22 | 0.37 | 94 | 1.33 | 1.67 | 1.83 |
| | 17 | M | 24 | 0.25 | 103 | 2.50 | 4.17 | 1.67 |
| | 18 | M | 23 | 0.70 | 119 | 4.17 | 3.33 | 5.00 |
| | 19 | F | 47 | 0.09 | 85 | 2.50 | 3.00 | 4.00 |
| | 20 | F | 22 | 0.43 | 133 | 3.33 | 5.00 | 6.67 |
| | 21 | F | 40 | 0.59 | 120 | 4.17 | 5.00 | 3.33 |
| | 22 | F | 24 | 0.54 | 124 | 1.67 | 4.17 | 2.50 |
| | 23 | M | 24 | 0.38 | 114 | 2.50 | 2.50 | 4.17 |
| | 24 | F | 18 | 0.26 | 128 | 1.67 | 4.00 | 4.17 |
| | 25 | F | 23 | 0.31 | 121 | 1.67 | 1.67 | 2.50 |
| | 26 | F | 30 | 0.92 | 97 | 6.67 | 5.00 | 9.17 |
| | 27 | F | 20 | 0.38 | 104 | 4.17 | 3.83 | 4.17 |
| | 28 | M | 23 | 0.60 | 125 | 0.00 | 3.33 | 0.00 |
| | 29 | F | 21 | 0.32 | 111 | 3.33 | 4.17 | 2.50 |
| | 30 | F | 22 | 0.23 | 87 | 4.00 | 5.00 | 5.00 |

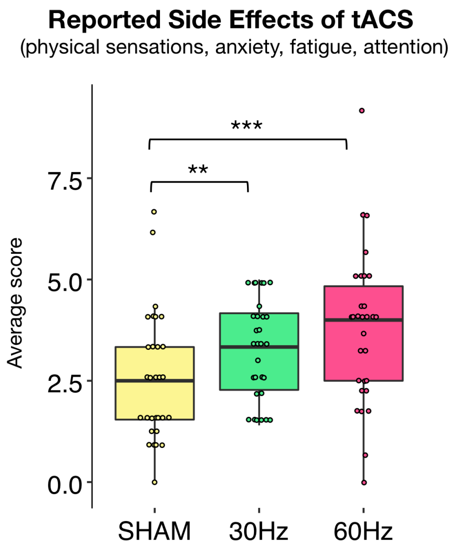


**Fig. S5. Demographic information and reported side effects of tACS.** Individual demographic information, language skills measured during the inclusion day (ECLA16+), non-verbal intelligence (Raven matrix) and reported side effects of tACS for each stimulation condition (sham/30 Hz/60 Hz). This included physical sensations - such as pain, warming, tingling – anxiety, fatigue and attention. The reported side effects were stronger for the active conditions as compared to the sham stimulation (** for $p < 0.01$, *** for $p < 0.001$). There was no difference between the dyslexia and no dyslexia group.

| Test | Example | Measured skills | | Items/ duration |
|---|---|---|---|---|
| **Pseudo-word repetition** | *Repeat:* DRONDOLINTIERBAPACHO | Phonological awareness (syllable and phoneme levels) | | 30 |
| | | -> Phoneme awareness | -> Syllabic short term memory | |
| **Spoonerism** | *Presented words* : **P**ATERE - **B**OUDIN<br>*Correct answer* : **B**ATERE - **P**OUDIN | Phonological awareness (word, syllable and phoneme level) | | 30 |
| | | -> Phoneme awareness | -> Verbal short term memory | |
| **Text reading** | "Certaines études ont mis l'accent sur le lien étroit entre l'émotion et la prise de décision. Il y a donc au moins deux processus distincts de prise de décision. L'un, basé sur une intelligence logicomathématique [..]" | Grapheme to phoneme conversion and global reading skills | | 3 min |
| | | -> Reading accuracy | -> Reading speed | |

**Table S1. Battery of linguistic tests**. Three linguistic tests were designed by a certified linguist to probe phonological processing, short-term memory and reading skills (speed and accuracy). The pseudo-word repetition test consisted in repeating non-lexical words that contain existing syllables in French. The spoonerism test consisted in transposing the first phoneme of two words, chosen to have similar phonological and syntactic features and with the same lexical frequency across sessions. Text reading consisted in reading a scientific text for 3 minutes.

| | DYSLEXIA (n=15) | | NO DYSLEXIA (n=15) | |
|---|---|---|---|---|
| | Mean | SD | Mean | SD |
| Age (Years) | 27.3 | 8.82 | 25.5 | 7.85 |
| Non-verbal IQ (APM) | 101.7 | 21.3 | 111 | 15.2 |
| **ECLA 16+ (Mean Z-Scores)** | | | | |
| Phonological awareness*** | -0.60 | 0.59 | 0.44 | 0.53 |
| Reading Skills*** | -0.65 | 0.9 | 0.28 | 0.52 |
| Short-term memory*** | -0.31 | 0.49 | 0.70 | 0.24 |
| **RAN (MeanZ-Scores)** | | | | |
| Rapid Automatized Naming (RAN)*** | -0.72 | 0.90 | 0.72 | 0.38 |

**Table S2. Dyslexia diagnosis session during the inclusion day.** Scores in the dyslexia and no dyslexia group for the ECLA 16+ diagnosis test and the Rapid Automatized Naming (RAN) test evaluating phonological awareness, reading skills and short-term memory. Negative values represent performance below average, positive values performance above average; all values are z-scores. Stars indicates significant differences between the dyslexia and no dyslexia group (*** for $p < 0.001$).
